# Control of mRNA decapping by autoinhibition

**DOI:** 10.1101/266718

**Authors:** David R Paquette, Ryan W Tibble, Tristan S Daifuku, John D Gross

## Abstract

5’ mediated cytoplasmic RNA decay is a conserved cellular process in eukaryotes. While the functions of the structured core domains in this pathway are understood, the role of abundant intrinsically disordered regions (IDRs) is lacking. Here we reconstitute the Dcp1:Dcp2 complex containing a portion of the disordered C-terminus and show its activity is autoinhibited by linear interaction motifs. Enhancers of decapping (Edc) 1 and 3 cooperate to activate decapping by different mechanisms: Edc3 alleviates auto-inhibition by binding IDRs and destabilizing an inactive form of the enzyme, whereas Edc1 stabilizes the transition state for catalysis. Both activators are required to fully stimulate an autoinhibited Dcp1:Dcp2 as Edc1 alone cannot overcome the decrease in activity attributed to the C-terminal extension. Our data provide a mechanistic framework for combinatorial control of decapping by protein cofactors, a principle that is likely conserved in multiple 5’ mRNA decay pathways.

---

Eukaryotic 5’-3’ mRNA decay is preceded and permitted by removal of the m^7^GpppN (N any nucleotide) cap structure and is a critical, conserved cellular process from yeast to humans (1–3). There are multiple decapping enzymes in eukaryotic cells (4), and the maintenance of the m^7^G cap in eukaryotes is required for numerous cellular processes including: splicing (5, 6), nuclear export (7), canonical translation and transcript stability (8, 9), and quality control pathways (10, 11). Recent evidence suggests that caps containing m^6^Am as the first-transcribed nucleotide are protected from decapping and 5’-3’ decay (12). Critically, mRNA decay is important during development (13). Furthermore, the cap structure and polyA tail differentiate the mRNA from other cellular RNAs, a feature viruses try to coopt to protect and translate their messages (14). Hydrolysis of the cap structure usually marks the transcript for degradation. Consequently, decapping and 5’-3’ decay pathways are tightly controlled and rely on cofactors (15). A key question in the field is how the activity of decapping enzymes are controlled by different protein interaction networks.

The major cytoplasmic decapping enzyme in yeast is the Dcp1:Dcp2 holoenzyme. Dcp2 is a bi-lobed enzyme consisting of a regulatory (NRD) and NUDIX containing catalytic domain (CD). The NRD binds Dcp1, an EVH1-like scaffold, which recruits cofactors through an aromatic cleft that recognizes a short proline-rich motif found in Edc1-type coactivators (16), including its yeast paralog Edc2 and mammalian PNRC2 (17). Additionally, Edc1-type coactivators contain a conserved short linear (Dcp2) activation motif that stimulates the catalytic step of decapping. Crystallographic and solution NMR analyses of Dcp1:Dcp2 with m7GDP product and substrate analog reveal the activation motif stabilizes the regulatory and catalytic domains of the enzyme in orientations compatible with formation of a composite active site (18; Mugridge et al., submitted). In the absence of Edc1 or related cofactors, Dcp1:Dcp2 is dynamic and forms nonproductive interactions with substrate RNA (18), which reduce catalytic efficiency. In several crystal structures of Dcp1:Dcp2, the composite active site is occluded (PDBID: 2QKM, 5J3Y, 5QK1), and it has been suggested that these forms of the enzyme exist in solution as nonproductive states of unknown function (18–20).

Edc1 has genetic interactions with Edc3 in yeast and both proteins form physical interactions with Dcp1:Dcp2 (21), suggesting they work together to promote decapping. Edc3 is important for decapping and subsequent 5’-3’ decay of pre-mRNA and mRNA targets in budding yeast (22, 23), general decapping in fission yeast (24), and decay of miRNA targets in *Drosophila* (25); while mutations in human Edc3 are associated with defects in neuronal development (26). Edc3 binds to fungal Dcp2 or metazoan Dcp1 through interactions with C-terminal extensions containing short leucine rich helical motifs (HLMs) (27). Recent crystallographic and genetic experiments in *S. cerevisiae* determined Pat1, a central component of the decapping machinery, also interacts with HLMs in Dcp2, demonstrating the importance of HLMs in mediating protein-protein interactions to control decay (28). Moreover, the HLMs are important for recruitment to P-bodies *in vivo* (20) and for phase-separation *in vitro* (27). However, the molecular mechanisms for how Edc3 stimulates decapping are not well understood.

Genetic studies in budding yeast indicate the disordered C-terminus of Dcp2 is a major site of regulation of decapping. He and Jacobson identified regions in the C-terminus that recruit positive regulators of decapping, including Edc3 and Pat1, to promote turnover of specific transcripts (29). The same study also demonstrated that a distinct proline- and phenylalanine-rich region in the C-terminus negatively regulates decapping and excision of this region bypasses the requirement for activation of decapping by Edc3 (29). These results suggest Dcp2 is autoinhibited, but a biochemical and structural understanding of how the C-terminus acts to inhibit decapping and how activators of decapping alleviate this inhibition is unknown due to the difficulty in preparing constructs of Dcp2 containing the disordered region.

Here we reconstitute an extended construct of *S. pombe* Dcp1:Dcp2 from recombinant components and show that it is autoinhibited by its C-terminal extension. Within this extension, we identify a proline-rich region that contains two inhibitory motifs that when removed restore enzymatic activity. We demonstrate that the addition of Edc3 alleviates the inhibitory role of the C-terminal extension by stimulating the catalytic step of decapping. Edc3 also promotes substrate binding. We show that a fraction of the autoinhibited complex is recalcitrant to Edc1 activation and that Edc3 works synergistically to make the complex more amenable to Edc1 dependent activation. Finally, we identify a conserved amino acid in the structured core domain of Dcp2 which when mutated quenches ms-µs dynamics of Dcp2, restores activity of the inhibited complex, and bypasses the Edc3 mediated alleviation of inhibition. We propose a model for autoinhibition of the decapping complex that occurs through stabilization of a cap-occluded Dcp2 conformation, which exists in solution and has appeared in numerous crystal structures.

## MATERIALS & METHODS

### Protein expression and purification

A pRSF containing polycistronic His-Gb1-tev-Dcp1:Dcp2(1-504)-strepII was used to coexpress the c-terminally extended Dcp1:Dcp2 complexes. Both the Dcp1 and Dcp2 sequences were codon-optimized for Escherichia coli from Integrated DNA Technologies and were cloned into a pRSF vector using Gibson assembly. The Dcp2 sequence contained a ribosome binding site (rbs) upstream and the Dcp1 was cloned behind the endogenous T7 promotor and rbs within the vector. The His-Gb1-tev-Dcp1:Dcp2(1-504)-strepII were expressed in E. coli BL21(DE3) (New England Biolabs) grown in LB medium. Cells were grown at 37 °C until they reached an OD_600_ = 0.6-0.8, when they were transferred to 4 °C for 30 minutes before induction by addition of 0.5mM IPTG (final concentration). Cells were induced for 16-18 hours at 18 °C. Cells were harvested at 5,000*g*, lysed by sonication (50% duty cycle, 3x1min), and clarified at 16,000*g* in lysis buffer (buffer composition goes here). The protein complex was purified in a two-step affinity purification: Ni-NTA agarose affinity column followed by strep-tactin high-capacity superflow and elution with 5mM d-desthiobiotin. The His-Gb1 tag was removed by addition of TEV protease overnight at 4 °C. The complex was further purified by size-exclusion chromatography on a GE Superdex 200 16/60 column in storage buffer (50mM HEPES, 100mM NaCl, 5% glycerol, 5mM DTT pH 7.5). The purified complex was concentrated, 20% v/v (final) glycerol was added, and then flash frozen in LN2 for kinetics studies. Dcp1:Dcp2(1-504) internal deletion constructs (IM1, IM2 and IM1&IM2) were generated by whole-plasmid PCR with 5’Phosphorylated primers. Dcp1:Dcp2 (1-504) Y220G was generated using whole-plasmid PCR with mutagenic divergent primers. A His-TRX-tev-Edc3 containing plasmid was generated by Gibson cloning S. pombe Edc3 cDNA into a pET30b plasmid which already contained a His-TRX-tev coding sequence. The LSm and YjeF N domain containing plasmids were created by whole-plasmid PCR with 5’Phosphorylated primers. The Edc3 constructs were purified as described (29) with a modification to the size-exclusion chromatography; storage buffer was used instead of the phosphate buffer as described.

### Kinetic Assays

Single-turnover *in vitro* decapping assays were carried out as previously described (31). Slight alteration to the buffer (brought up volume in storage buffer with additional 20% v/v glycerol). A ^32^P-labeled 355-nucleotide RNA substrate (containing a 15-nucleotide poly(A) tail) derived from *S. cerevisae* MFA2 mRNA was used for all the decapping assays; where the m^7^G cap is radiolabeled on the α phosphate such that, upon decapping, the excised m^7^GDP product could be detected by TLC and autoradiography. Reactions were initiated by mixture of 30 μL 3× protein solution with 60 μL 1.5× RNA solution at 4 °C; final Dcp1:Dcp2 concentration was 1.5μM and the final RNA concentration was <100 pM. For decapping assays containing Edc3; Edc3 was added in 4-fold molar excess and the mixture was incubated at RT for 20 minutes before transferring to the 4 °C block. For decapping assays containing Edc1, the peptide (synthesized by Peptide2.0) was added at the indicated concentration and the mixture was incubated at RT for 20 minutes before transferring to the 4 °C block. Samples were equilibrated for at least 30 minutes on the 4 °C block before initiating. Time points were quenched by addition of excess EDTA, TLC was used to separate the RNA from the product m^7^GDP, and the fraction decapped was quantified with a GE Typhoon scanner and ImageQuant software. Fraction m^7^GDP versus time were plotted and fit to a 1st-order exponential to obtain k_obs_; in the case of Dcp1:Dcp2(1-504) when the kinetics were too slow to obtain reliable exponential fits, k_obs_ was obtained from a linear fit of the initial rates by division of the slope by the empirically derived endpoint.

### NMR Spectroscopy

ILVMA methyl labeling of Dcp2 was carried out in D_2_O M9 minimal media with ^15^NH_4_Cl and ^12^C/^2^H-glucose as the sole nitrogen and carbon sources, respectively, and labeled precursors (Ile: 50 mg L^-1^, Leu/Val: 100 mg L^-1^, Met: 250 mg L^-1^, Ala: 100 mg L^-1^) were added 40 minutes prior to induction. Following overnight induction with 1 mM IPTG, cells were lysed by sonication and clarified at 16,000*g*. Dcp2 was purified by incubating clarified lysate with nickel resin followed by elution with 250 mM imidazole. The His-GB1-TEV tag was then removed by digestion with TEV protease and untagged Dcp2 was run over a Superdex 75 size exclusion chromatography column equilibrated with pH 7.0 NMR buffer (21.1 mM NaH_2_PO_4_, 28.8 mM Na_2_PO_4_, 200 mM NaCl, 100 mM Na_2_SO_4_, 5 mM DTT). All ^1^H-^15^N HSQC and CPMG experiments were performed with 250 μM protein at 303K on a Bruker Avance 800 spectrometer equipped with a cryogenic probe.

For CPMG analysis, dispersion curves were acquired with a 40-ms constant-time delay wherein the pulse frequency was varied between 50 and 950 Hz. FuDA (gift from D.F. Hansen, University College London, London, UK, http://www.biochem.ucl.ac.uk/hansen/fuda/) was used to extract intensities, which were converted to relaxation rates using procedures outlined in (32). Errors in R_2,eff_ are reported as the pooled standard deviation determined using procedures outlined in (33). Dispersion curves were fit to a two-site exchange model using the program cpmg_fitd8 (gift from D. Korzhnev and L. Kay, University of Toronto, Toronto, ON).

## RESULTS

### A segment of the C-terminus in Dcp2 inhibits decapping

The C-terminus of *S. pombe* Dcp2 is predicted to be highly disordered (**Fig 1A**). To determine its functional role in decapping, Dcp1 was co-expressed with the C-terminally extended Dcp2 (Dcp1:Dcp2_ext_) and purified to homogeneity. A C-terminal boundary ending at residue 504 was chosen by sequence conservation and optimization of expression. This construct contains regions of the protein that were excised in prior biochemical studies due to sub-optimal expression levels (29, 32). Decapping activity of Dcp1:Dcp2_ext_ was compared with that containing only the structured core domains (Dcp1:Dcp2_core_) comprised of Dcp1 and Dcp2 with the N-terminal regulatory (NRD) and catalytic domain (CD) using a cap-radiolabeled RNA as substrate. The rate of decapping by Dcp1:Dcp2_core_ was faster than Dcp1:Dcp2_ext_ (**Fig 1B**). Fitted rate-constants indicated Dcp1:Dcp2_ext_ was consistently slower than Dcp1:Dcp2_core_ (**Fig 1C,D**); this effect was not dependent on enzyme preparation. Dcp1:Dcp2_ext_ purified as a well-resolved, homogeneous peak on gel-filtration and is monodisperse under decapping reaction conditions, as indicated by dynamic light-scattering (**Fig S1A,B**). These data suggest the C-terminal extension of Dcp2 has sequence motifs that can inhibit the decapping activity of the structured, core domains.

**Figure 1:**
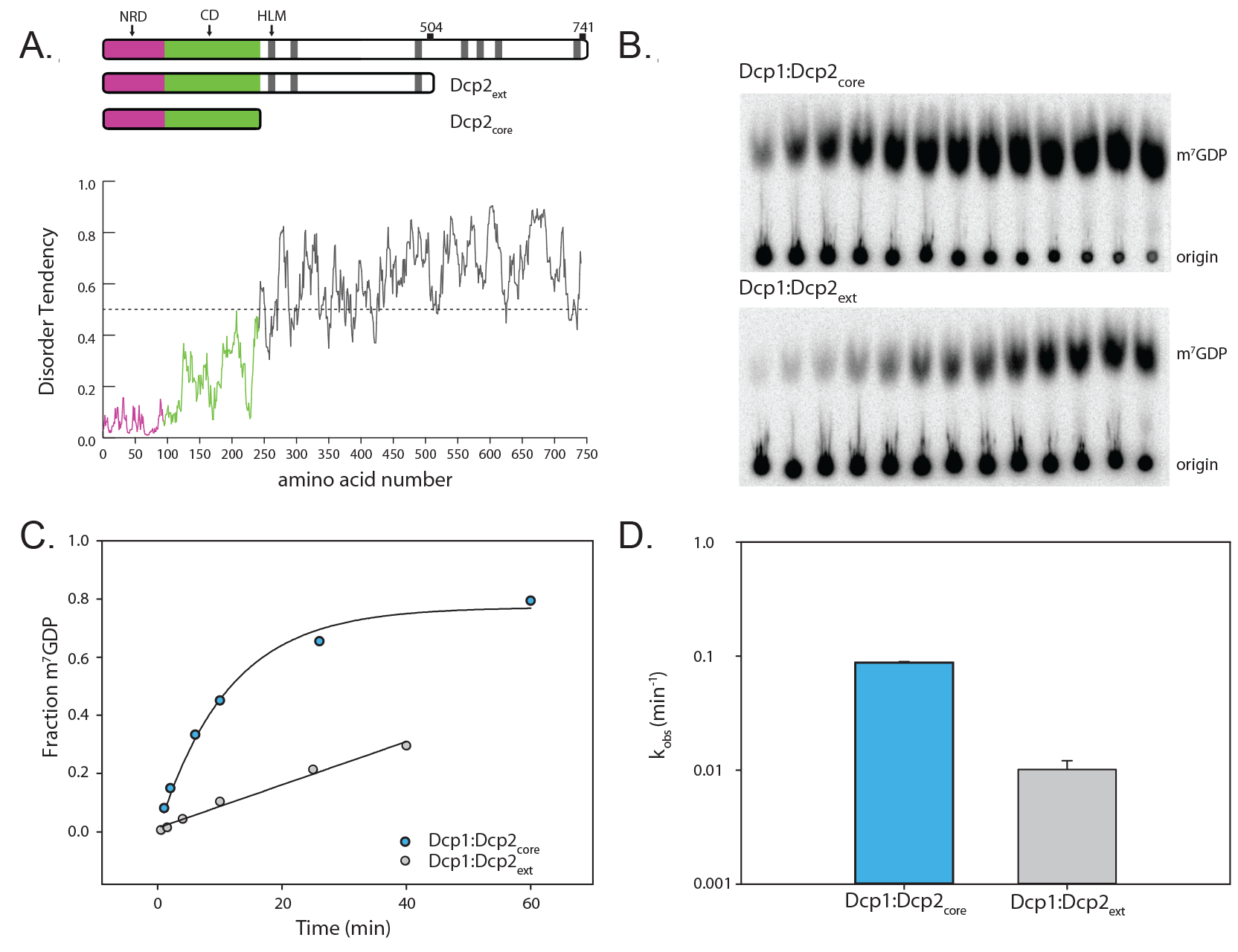
The C-terminal extension of Dcp2 inhibits decapping **(A)** Block diagram of the domains of S. pombe Dcp2. The magenta box (1-94) comprises the N-terminal regulatory domain (NRD) and the green (95-243) comprises the catalytic domain, which contains the Nudix helix. Grey bars are helical leucine-rich motifs (HLMs). The disorder tendency plot below was calculated using the IUPRED server (52). Where regions above the dotted line have a higher propensity for being disordered. (**B**) Representative raw TLC (thin-layer chromatography) showing the formation of m7GDP product over 40 minutes. The lower spots are the RNA origin and the upper are the cleaved cap. (**C**) Representative plot of fraction m7GDP product versus time comparing the catalytic core (Dcp1:Dcp2_core_) and inhibited C-terminally extended Dcp1:Dcp2_ext_. (**D**) A log-scale plot of the empirically determined rates from (C), where the error bars are the population standard deviation, σ. Difference in measured rates are significant as determined by unpaired t-test (see Table S3).

### Two motifs are required for autoinhibition of Dcp1:Dcp2

Having determined that Dcp1:Dcp2_ext_ was less active than Dcp1:Dcp2_core_, we next asked which regions in the C-terminal extension confer inhibition. Since the C-terminal extension of Dcp2 is predicted to be disordered, we hypothesized that inhibition of decapping may be mediated by linear interaction motifs. Candidate inhibitory motifs were queried by analysis of sequence conservation amongst the most closely related fission yeast, as linear motifs are known to evolve rapidly due to a lack of restraints imposed by three-dimensional structure (35, 36). Using this approach, we identified two possible inhibitory motifs: a proline-rich sequence (PRS) that aligns to the inhibitory element identified in budding yeast (29) previously shown to bind fission yeast Dcp1 (37), and a highly conserved stretch of amino acids that strongly resembles Dcp1-binding motifs in Edc1-like coactivators (**Fig 2A,S2**). We term these regions IM1 and IM2, respectively. Next, we queried whether deletion of these conserved regions, alone or in combination, would alleviate autoinhibition. The Dcp1:Dcp2 complexes where either putative IM1 or IM2 were deleted alone or in tandem were purified to homogeneity (**Fig S1**). Deletion of a region containing IM1 (residues 267-350 in Dcp2) revealed that it partially contributes to autoinhibition in Dcp1:Dcp2_ext_ (**Fig 2B,C**); this region was previously observed to have no effect on decapping activity when added in *trans* to Dcp1:Dcp2_core_ (37). Likewise, deletion of IM2 restored activity in Dcp1:Dcp2_ext_ to a similar degree as deletion of IM1 (**Fig 2B,C**). Furthermore, we found that a peptide of IM2 (residues 399-432) directly interacts with Dcp1:Dcp2_core_ in *trans* (**Fig S2C**). The combined removal of these regions fully restored the activity of Dcp1:Dcp2_ext_ to that of Dcp1:Dcp2_core_ (**Fig 2B,C**). Additionally, internal deletions of nonconserved regions in the C-terminal extension did not show an increase in activity relative to Dcp1:Dcp2_ext_ (**Fig S2D**). We conclude that the two linear motifs we identified in the C-terminal extension of fission yeast Dcp2, IM1 and IM2, are responsible for the autoinhibition of Dcp1:Dcp2.

**Figure 2:**
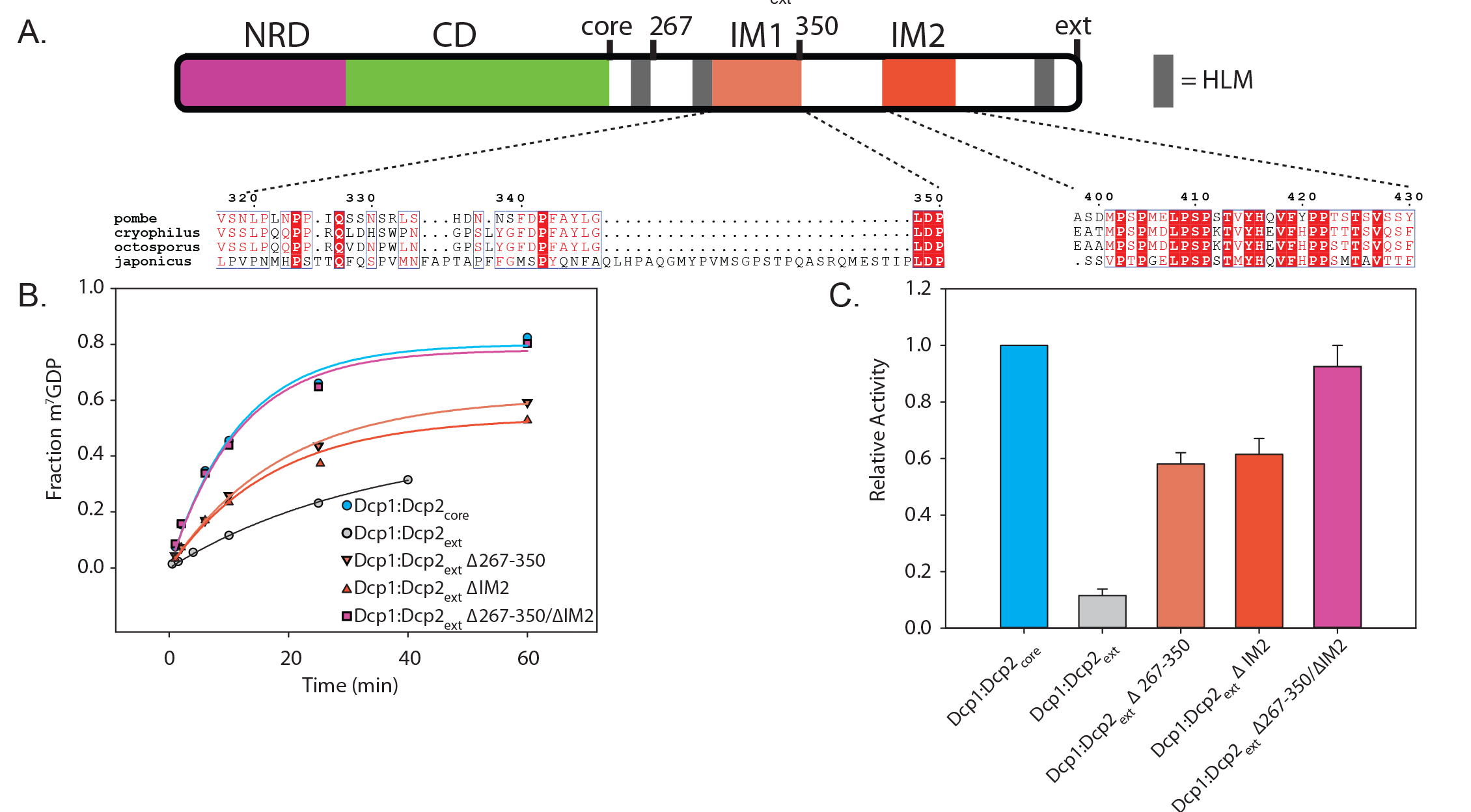
Two motifs are required for autoinhibition of Dcp1:Dcp2_ext_. (**A**) Block diagram colored as in Fig1. with IM1 and IM2 regions colored and the sequence conservation (53) for each motif shown below. IM1 contains proline and phenylalanine residues similar to the negative regulatory element identified in budding yeast (Fig S2B), while IM2 is absolutely conserved in all fission yeast. (**B**) Plot with fits for fraction of m7GDP versus time comparing the activity of Dcp1:Dcp2_ext_ where either IM1, IM2 or both are internally deleted. (**C**) Bar graph of the relative enzymatic activity of the various Dcp1:Dcp2_ext_ complexes compared to Dcp1:Dcp2_core_. Each IM contributes to the inhibitory effect of the c-terminal regulatory region (CRR). The error bars are the population standard deviation, σ. Differences in observed rates are significant except for Dcp1:Dcp2_core_ relative to Dcp1:Dcp2(Δ267-350/ΔIM2) and Dcp1:Dcp2(Δ267-350) relative to Dcp1:Dcp2(ΔIM2) as determined by unpaired t-test (see Table S3).

### A conserved surface on the catalytic domain of Dcp2 is required for autoinhibition

Enzymes that are regulated by autoinhibition typically exist in an inactive conformation that is distinct from the catalytically active form, which can block substrate recognition and catalysis (38). Typically, this entails linear interaction motifs interacting with core domains, which maintains the enzyme in the inactive state. We and others have suggested that the ATP-bound, closed Dcp1:Dcp2 structure (Song et al 2008, PDB 2QKM) could resemble an inactive conformation of the enzyme since the substrate binding site is occluded (20, 39). Recently, it was shown by NMR that this closed form of Dcp1:Dcp2 is significantly populated in solution (18). We hypothesized the autoinhibited form of Dcp1:Dcp2 might correspond to this cap-occluded conformation. This conformation is stabilized by several conserved residues that anchor the NRD to the CD of Dcp2, including W43, D47 and Y220 (Fig S3). In this conformation, Y220 blocks access of W43 and D47 of the NRD to cap. When Dcp2 is bound to m7G of cap in the catalytically-active conformation, Y220 is displaced and W43 and D47 of the NRD make essential interactions with m^7^G (**Fig 3A**) (37, 40; Mugridge et al. submitted). Therefore, we reasoned mutation of Y220 would destabilize the cap-occluded form of Dcp1:Dcp2, permitting it to more readily populate the cap-accessible, active conformation. Mutation of Y220 to glycine enhanced decapping activity of the Dcp1:Dcp2_core_ by 1.5-fold (Fig 3B) and we were able to observe cap binding in Dcp2_core_ by NMR (**Fig S4**). We next assessed the effect of the Y220G mutation in Dcp1:Dcp2_ext_ and observed a 6-fold increase in activity relative to wild-type Dcp1:Dcp2_ext_ (**Fig 3B**). These data suggest the cap-occluded conformation of Dcp1:Dcp2, which is observed in solution and in many crystal structures, could resemble the autoinhibited form of the enzyme containing the C-terminal extension.

**Figure 3:**
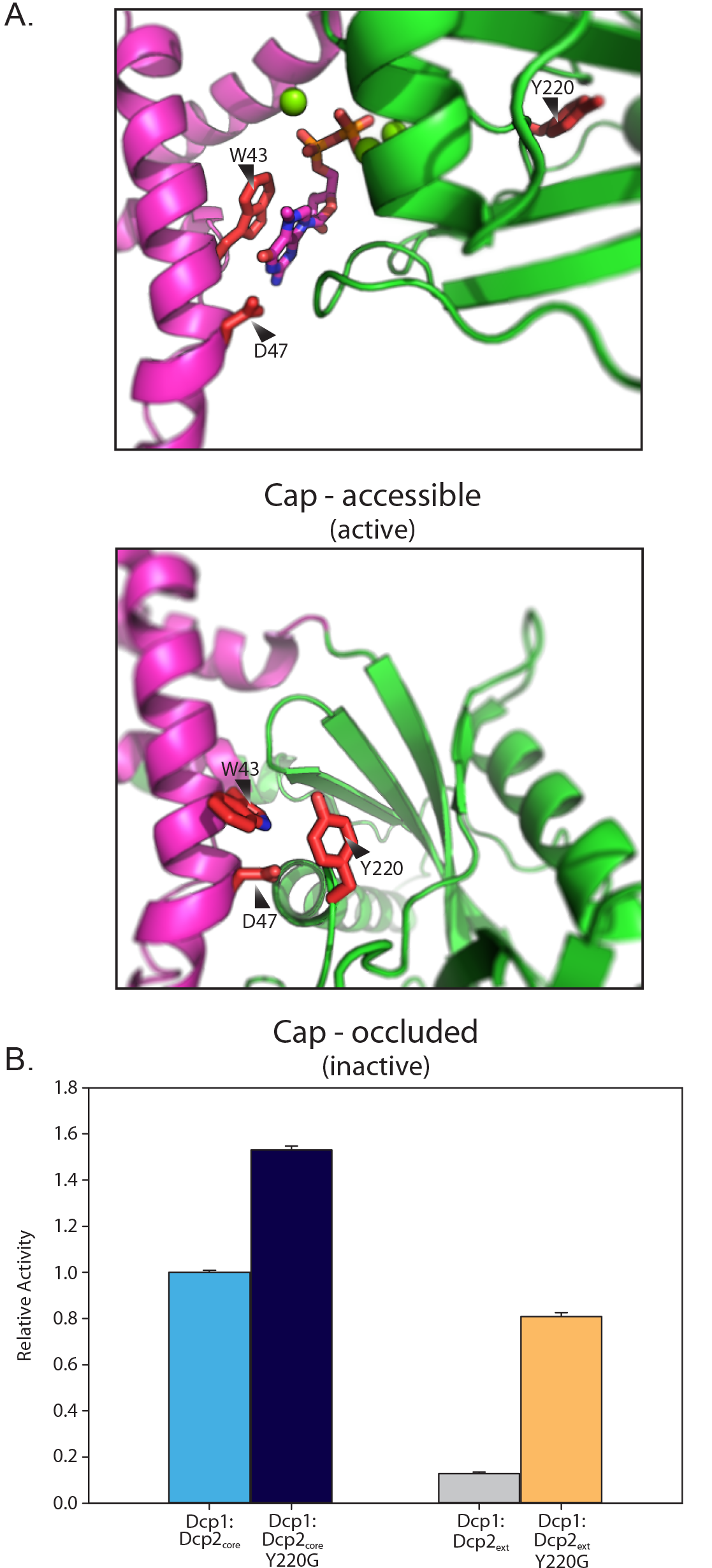
Y220 stabilizes a cap-occluded state and alleviates inhibition. (**A**) W43 and D47 that coordinate the m7G cap exist in conformations where they are either accessible or occluded by interaction with the conserved Y220G. (**B**) Plot of the relative activity of WT or Y220G Dcp1:Dcp2_ext_compared to the same mutation in Dcp1:Dcp2_core_. Dcp1:Dcp2_ext_ exhibits a greater increase in k_obs_ when Y220 is mutated as determined by an *in vitro* decapping assay. The error bars are the population standard deviation, σ. All differences in reported rates are significant as determined by unpaired t-test (see Table S3).

The Y220G mutation may activate decapping by destabilizing the cap-occluded conformation. To test this possibility, we used dynamic NMR spectroscopy. Previously, it had been noted that Dcp2 experiences global motions on the millisecond to microsecond timescale (ms-μs dynamics) where the NRD and the CD sample open and closed, cap-occluded forms, with Dcp1 enhancing the population of the latter state (18, 19). W43 was identified as a “gatekeeper” required for these dynamics (19). Since Y220 makes interactions with W43 in the cap-occluded state, we reasoned the Y220 mutation would also quench global ms-µs dynamics, resulting in the reappearance of broadened cross-peaks. Upon mutation of Y220 to glycine in Dcp2_core_, the expected cross-peaks from both the NRD and CD reappeared (**Fig S4**), indicating dynamics were quenched similarly to the previously reported W43 mutation (**Fig 4A**)(19). To corroborate that mutation of Y220 quenched the collective dynamics, we used CPMG spectroscopy on ^13^C-methyl ILVMA side-chain labeled Dcp2. Consistent with previous experiments, significant dephasing of transverse magnetization from collective ms-µs motions in wild-type Dcp2 could be attenuated with increasing CPMG pulse rate (**Fig 4B, black filled circles**) (18,19). In contrast to the wild-type, these dynamics are not present for the Y220G mutant (**Fig 4B, blue circles**). We conclude that the Y220G mutation shifts the equilibrium from cap-occluded to the open conformation.

**Figure 4:**
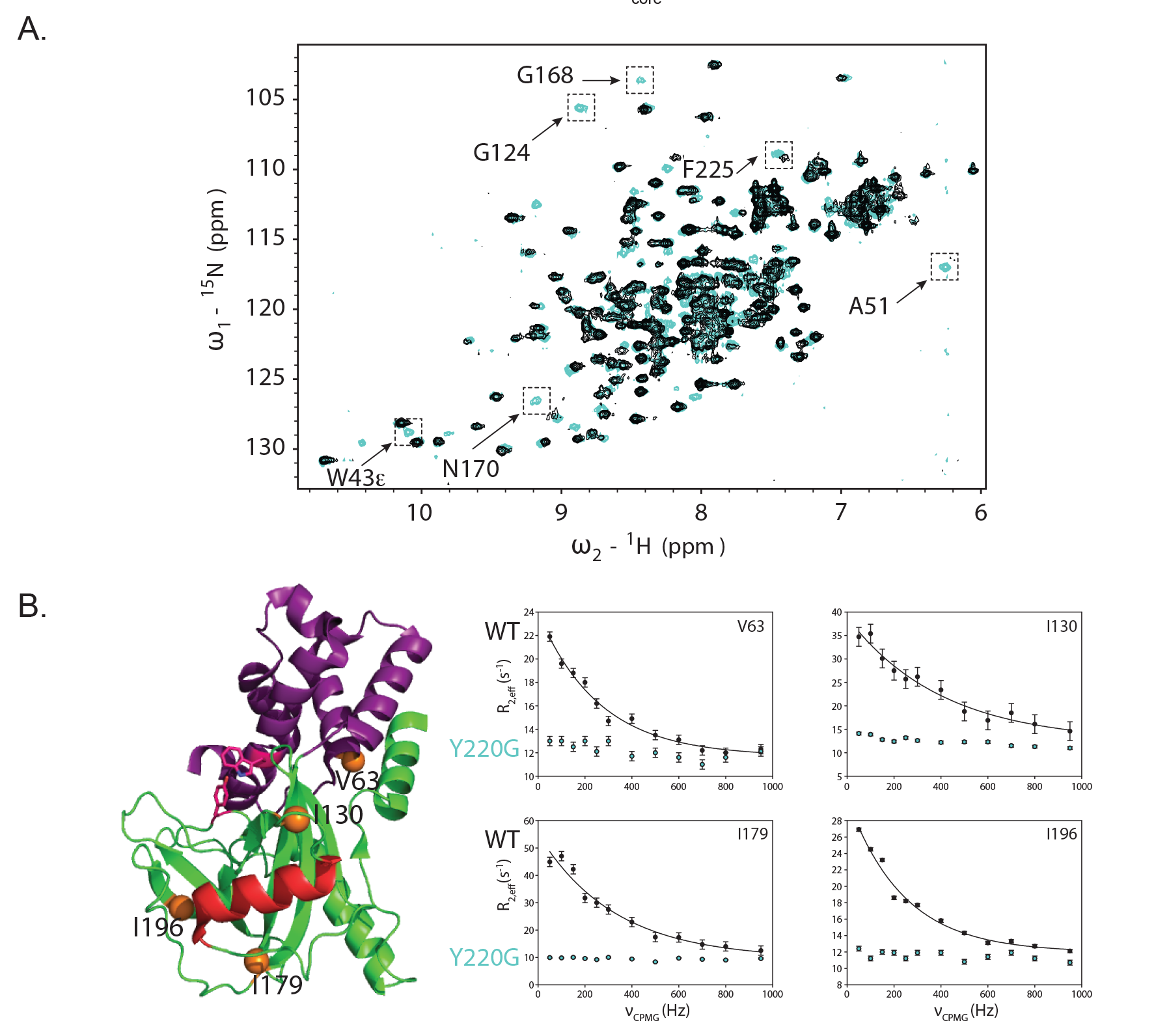
Y220G mutation quenches ms-μsec dynamics in Dcp2_core_. (**A**) Shown are the ^15^N HSQC spectra of WT Dcp2 residues 1-243 (black) and Dcp2 Y220G (cyan). Selected residues with significant changes upon mutation are indicated. (**B**) Location of the four representative residues whose sidechain dynamics are presented are shown as orange spheres on the ATP-bound structure of DCP2 (2QKM), where the NRD is magenta, CD is green, catalytic Nudix helix is red and W43 and Y220 are displayed as sticks. Selected CPMG dispersion data at 800 MHz for WT (black circles) or Y220G (cyan circles). WT data fit well to a two-site exchange model (Black lines), whereas Y220G data did not, indicating ms-μs dynamics are strongly reduced.

### Edc3 alleviates autoinhibition of Dcp1:Dcp2_ext_ by stimulating the catalytic step

Since the inhibitory motifs of Dcp2 are proximal to known Edc3 binding sites (the HLMs) we tested if Edc3 could alleviate autoinhibition. When Dcp1:Dcp2_ext_ was incubated with saturating concentrations of Edc3, a stable complex was observed and decapping activity was completely restored to rates observed for Dcp1:Dcp2_core_ (**Fig 5A,B; Fig S5; Table S2**). Having identified a mutation in the catalytic domain of Dcp2_core_ that destabilizes autoinhibition, we next examined whether this mutation bypasses activation by Edc3. Introduction of the Y220G mutation into Dcp1:Dcp2_ext_ markedly attenuates stimulation by Edc3 and closely resembles the extent of activation observed for Dcp1:Dcp2_HLM-1_, which is a construct of Dcp1:Dcp2_core_ containing a single Edc3 binding site (**Fig 5C**). The differential activation observed for Dcp1:Dcp2:Edc3 complexes with and without the C-terminal extension containing autoinhibitory motifs indicates Edc3 utilizes multiple mechanisms to enhance decapping.

**Figure 5:**
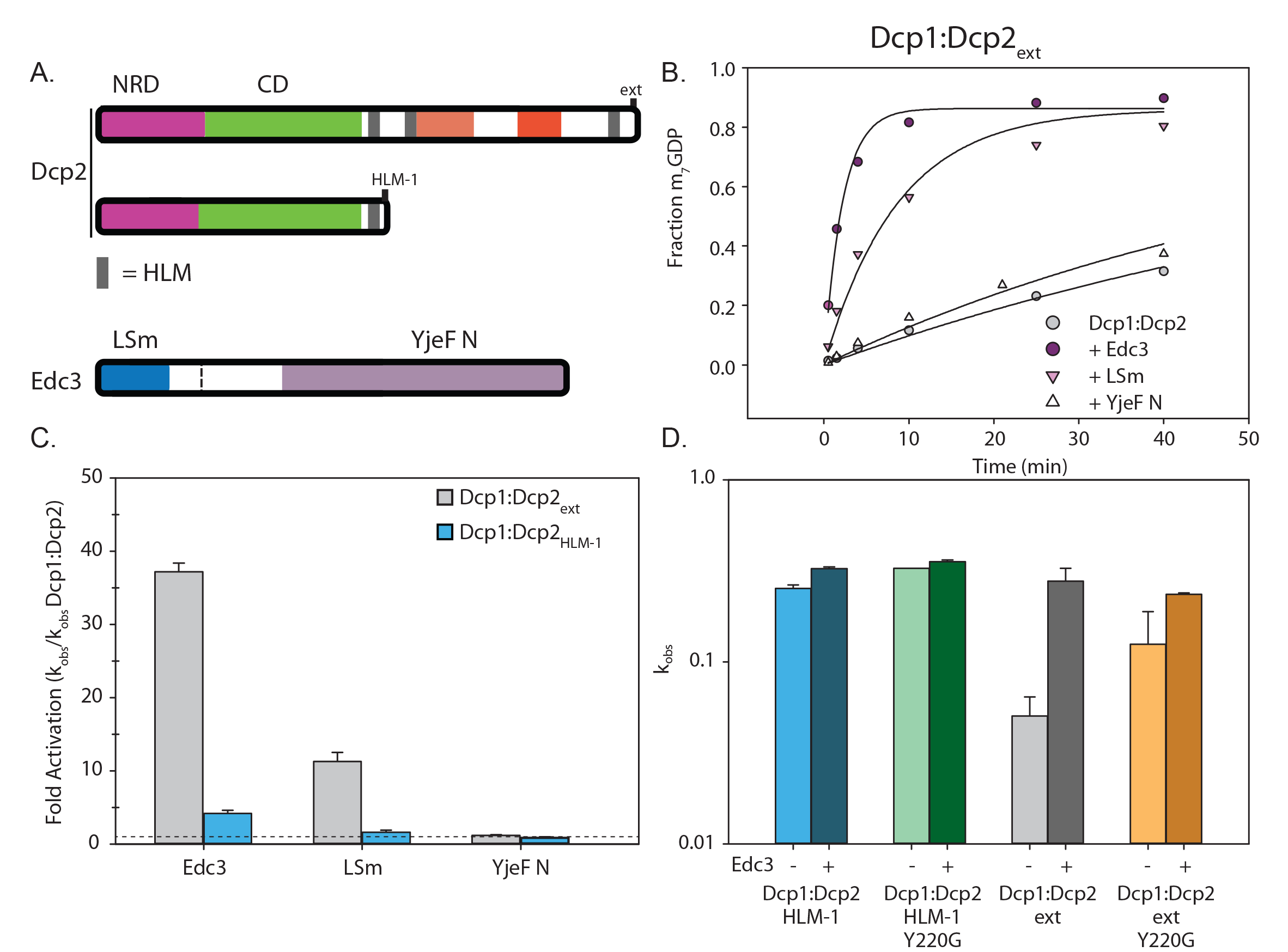
Edc3 alleviates autoinhibition of Dcp1:Dcp2_ext_. (**A**) Block diagram of the Dcp2 and Edc3 used in the subsequent decapping assay dc3 consists of an LSm domain that interacts with HLMs and a YjeF N domain that provides an RNA binding surface when dimerized. (**B**) Decapping activity of Dcp1:Dcp2_ext_ incubated with excess Edc3, LSm domain or YjeF N domain. (**C**) Comparison of the relative fold activation of Dcp1:Dcp2_ext_ versus Dcp1:Dcp2_HLM-1_ with the various Edc3 constructs. All relevant differences are significant as determined by unpaired t-test (see Table S3 for p-values for all pairwise comparisons). (**D**) Logscale plot of decapping rate of Dcp1:Dcp2_HLM-1_ [blue], Dcp1:Dcp2_HLM-1_ Y220G [green], Dcp1:Dcp2_ext_ [gray], or Dcp1:Dcp2_ext_ Y220G [yellow] where the darker bar is the rate with excess Edc3. The error bars are the population standard deviation, σ. Differences in observed rates are significant except for Dcp1:Dcp2_HLM-1_ Y220G relative to Dcp1:Dcp2_HLM-1_ Y220G:Edc3 and Dcp1:Dcp2_ext_ Y220G relative to Dcp1:Dcp2_ext_ Y220G:Edc3 as determined by unpaired t-test (see Table S3 for p-values of all pairwise comparisons).

Edc3 is known to interact with HLMs in the C-terminus of Dcp2 through its N-terminal LSm domain and has been shown to bind RNA through its C-terminal Yjef N domain (20, 41). To dissect the contributions from these domains in enhancing decapping, we compared their activation of Dcp1:Dcp2_HLM-1_ and Dcp1:Dcp2_ext_ (**Fig 5D, Fig S5B**). The Edc3 LSm domain was sufficient to increase the activity of Dcp1:Dcp2_ext_ to levels comparable with Dcp1:Dcp2_core_; whereas addition of the Edc3 Yjef N domain in isolation had no effect on decapping activity (**Fig 5D**). These results indicate binding of the Edc3 LSm domain to the HLMs in Dcp2 is sufficient to alleviate autoinhibition of the C-terminal regulatory region of Dcp2, but full-activation requires the Yjef N domain. In contrast to Dcp1:Dcp2_ext_, the LSm domain of Edc3 does not provide any substantial increase in activity of Dcp1:Dcp2_HLM-1_; whereas a 5-fold increase in decapping is observed upon addition of full-length Edc3. These observations suggest Edc3 activates decapping both by alleviating autoinhibition to enhance the catalytic step and by promoting RNA binding.

To test these possibilities, we measured rates of decapping by Dcp1:Dcp2 in the presence and absence of saturating Edc3 under single-turnover conditions. This approach allows us to determine the contributions of Edc3 activation to RNA binding (K_m_) and the catalytic step of decapping (k_max_). Compared to Dcp1:Dcp2_HLM-1_, Dcp1:Dcp2_ext_ had a significant 6-fold reduction in k_max_, indicating autoinhibition occurs in the catalytic step (**Fig 6A-D**, compare lower curves in **A,B**; light blue and light gray bars in **C,D**). When kinetics for Dcp1:Dcp2_ext_ were performed in the presence of Edc3, there was a substantial 6-fold increase in k_max_ and concomitant 5-fold decrease in K_m_ relative to Dcp1:Dcp2_ext_ alone (**Fig 6B,D**).

Thus, the observed 30-fold stimulation of Dcp1:Dcp2_ext_ by Edc3 (**Fig 5C,D**) can be broken down as 6-fold from alleviation of autoinhibition by the its LSm domain binding HLMs in the extended C-terminus of Dcp2, and 5-fold deriving from RNA binding by Edc3. The effects of Edc3 on Dcp1:Dcp2_ext_ activity are in contrast to Dcp1:Dcp2_HLM-1_, which in the presence of Edc3 had a similar 5-fold decrease in K_m_ but not a significant change in k_max_ (**Fig 6A,C**). The observed 5-fold decrease in K_m_ for both Dcp1:Dcp2_HLM-1_ and Dcp1:Dcp2_ext_ is in good agreement with the reported ability of Edc3 to bind RNA and suggests Edc3 predominantly provides additional RNA binding capacity in the absence of autoinhibition (40, 42). Therefore, the mechanism of activation by Edc3 is a combination of binding to HLMs in order to alleviate autoinhibition and enhance the catalytic step while providing an additional RNA binding surface to increase Dcp2 substrate affinity.

**Figure 6:**
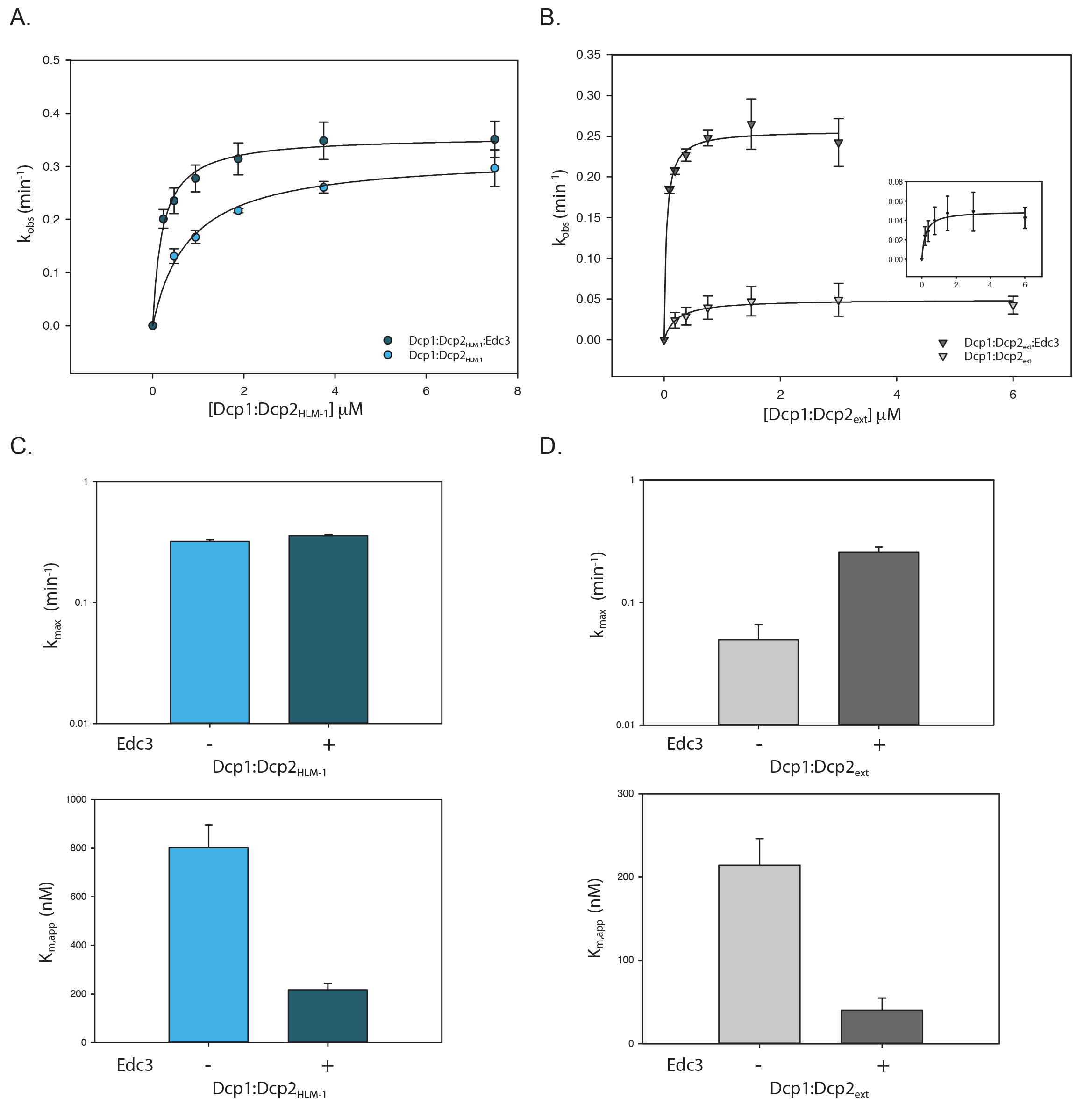
Edc3 alleviates autoinhibition and promotes RNA binding. (**A**) Plot of k_obs_ versus Dcp1:Dcp2_HLM-1_ concentration in the absence (light blue) or presence (dark blue) of saturating concentrations of Edc3. Error bars represent population standard deviation, σ. (**B**) Plot of k_obs_ versus Dcp1:Dcp2_ext_ concentration in the absence (light gray) or presence (dark gray) of saturating concentrations of Edc3. Error bars represent population standard deviation, σ. (**C**) Comparison of fitted k_max_ (top) and K_m,app_ (bottom) values from panel A for Dcp1:Dcp2_HLM-1_ with or without saturating Edc3 (colored as in panel A). There is not a significant change in k_max_ upon addition of Edc3 (as determined by unpaired t-test, see Table S3) but K_m,app_ is 5-fold decreased. (**D**) Comparison of fitted k_max_ (top) and K_m,app_ (bottom) values from panel B for Dcp1:Dcp2_ext_ with or without saturating Edc3 (colored as in panel B). There is a 6-fold increase in k_max_ upon incubation with Edc3 and K_m,app_ decreases by 5-fold. Error bars are the population standard deviation, σ.

### Edc3 and Edc1 work together to activate decapping by Dcp1:Dcp2_ext_

Edc1 stimulates the catalytic step of decapping by stabilizing the composite cap binding site of Dcp2 formed by the N-terminal regulatory and catalytic domains (18; Mugridge et al., submitted). Here we have shown Edc3 alleviates autoinhibition by binding sites distal from the Dcp1:Edc1 interaction site, which suggests these activators could work together to promote decapping. To test this possibility, we titrated a peptide of spEdc1 (residues 155-180) that contains the minimal Dcp1 binding and Dcp2 activation motifs (Edc1-DAM, see ref 43) against Dcp1:Dcp2_ext_ or Dcp1:Dcp2_core_ and determined if Edc3 changed the threshold of activation, defined as a con-Figure 7: Edc1 and Edc3 coordinate to activate the centration of Edc1-DAM required for one-half maximal activDcp1:Dcp2_ext_ complexity (K_1/2_) (**Fig 7A**). For experiments with Dcp1:Dcp2_ext_, the K_1/2_ was shifted 3.5-fold higher compared to the Dcp1:Dcp-2_core_ (**Fig 7B**). In addition, even at saturating concentrations, Edc1 was unable to fully overcome the inhibitory effect of the Dcp2 C-terminal extension in the absence of Edc3. In contrast, addition of Edc3 returned the K_1/2_ for Edc1-DAM activation of Dcp1:Dcp2_ext_ to within error of Dcp1:Dcp2_core_ and rescued the maximal observed rate. A maximum rate of 6.2 min^-1^ was observed when Edc1 and Edc3 were present at saturating concentrations, which is greater than the maximum rate observed for the Dcp1:Dcp2_core_ saturated with Edc1. These observations suggest Edc1-DAM alone is not sufficient to fully activate decapping of Dcp1:Dcp2_ext_ and instead requires Edc3 to fully stimulate the rate of the autoinhibited complex greater than 500-fold (**Fig S6**).

**Figure 7:**
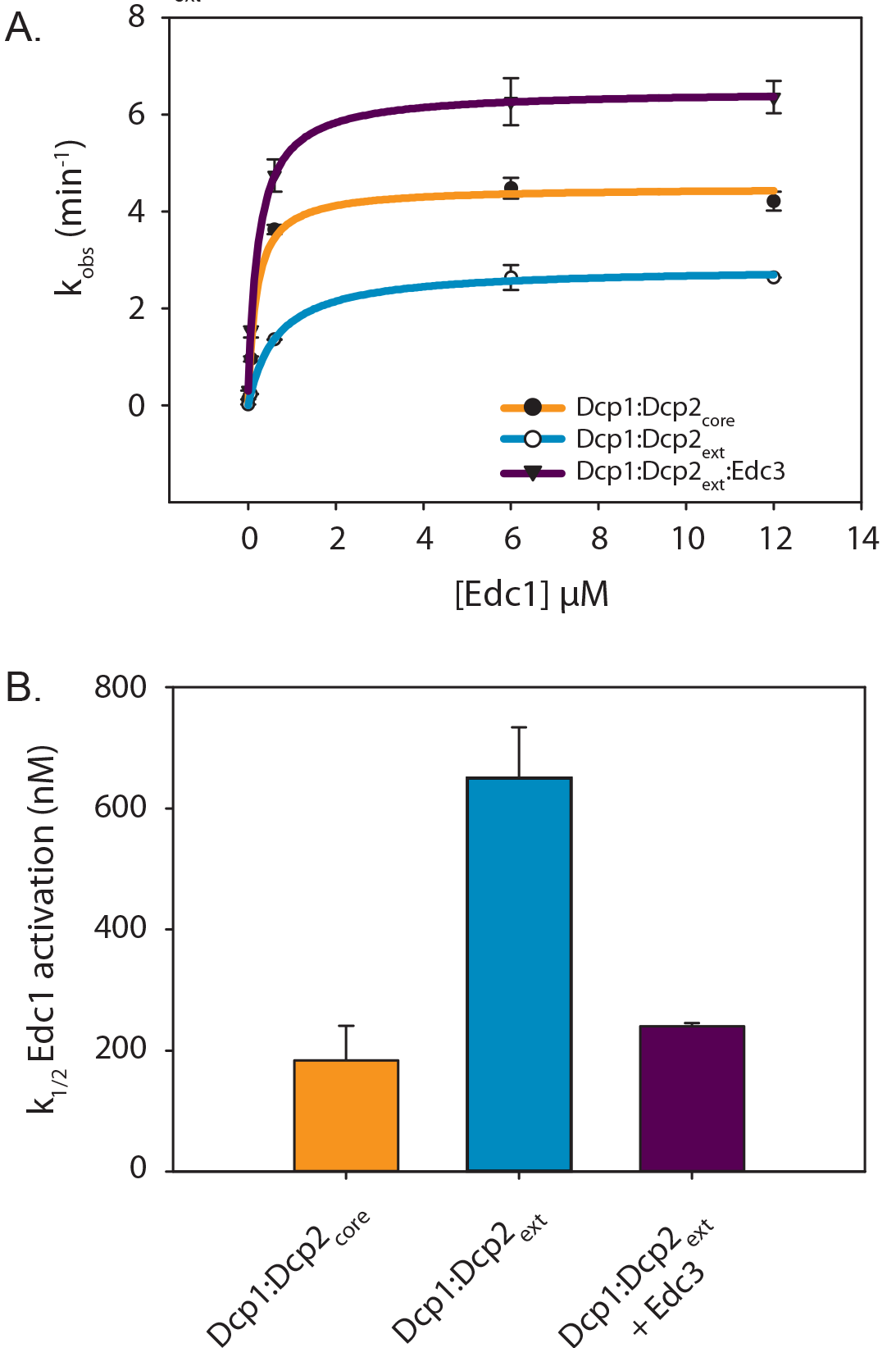
Edc1 and Edc3 coordinate to activate the Dcp1:Dcp2_ext_ complex. (**A**) Plot of k_obs_ versus Edc1 concentration for the catalytic core (orange), autoinhibited complex (cyan), or the autoinhibited complex (Dcp1:Dcp2_ext_) saturated with Edc3 (purple). Error bars are population standard deviation, σ. (**B**) Bar graphs showing the k1/2 Edc1 activation (apparent kd) determined from the fits in panel A for Edc1 dependent activation with same colors as in A. Error bars are standard error. Differences in k1/2 are significant except for Dcp1:Dcp2_core_ relative to Dcp1:Dcp2_ext_:Edc3 as determined by unpaired t-test (see Table S3).

## DISCUSSION

The structures of a majority of the globular domains of the mRNA decay proteins are available and have provided key insights into how these factors interact with one another to mediate decay (27). While progress has been made on this front, much less is known regarding the regulatory functions of intrinsically disordered regions (IDRs) that are replete in these proteins. Here, we have determined that the disordered C-terminal extension of fungal Dcp2 contains two motifs (IM1 and IM2) that inhibit decapping activity. The inhibitory effect of the disordered extension likely relies on an underlying cap-occluded conformation, which is stabilized by a conserved aromatic side-chain in the CD of Dcp2. Edc3 alleviates autoinhibition by binding the HLMs that are proximal to the inhibitory motifs and provides increased RNA binding affinity through its RNA binding domain. Stimulation of Dcp1:Dcp2_ext_ by Edc1 requires Edc3 to achieve maximal activation (∼500-fold), and we provide evidence that such control of decapping is achieved by factors acting on distinct conformational substates.

Decapping by Dcp1:Dcp2 requires formation of a composite active site shaped by residues lining the NRD and the CD of Dcp2. The core complex is dynamic in solution, sampling an open form where residues that line the composite active site are far apart, or a closed form where residues on the NRD that engage cap to promote catalysis are occluded (18). Several observations suggest the closed, cap-occluded form of Dcp1:Dcp2 is representative of the autoinhibited form containing the C-terminal extension (**Fig 8A**). First, this conformation has been observed in a variety of crystal structures and in solution (18, 40, 41, 43, 44). Second, it is incompatible with cap recognition, since essential residues of the NRD that contact the m^7^G moiety (W43 and D47) are blocked by a conserved amino acid on the CD (Y220) (**Fig S3**). Third, mutation of a conserved Tyr predicted to stabilize this nonproductive state increases the catalytic activity, destabilizes the “occluded” state in solution and disrupts the inhibitory effect of the Dcp2 extension (**Figs 3,4**). Fourth, the C-terminal extension reduces that catalytic step of decapping, consistent with the composite active site of Dcp2 being occluded.

**Figure 8:**
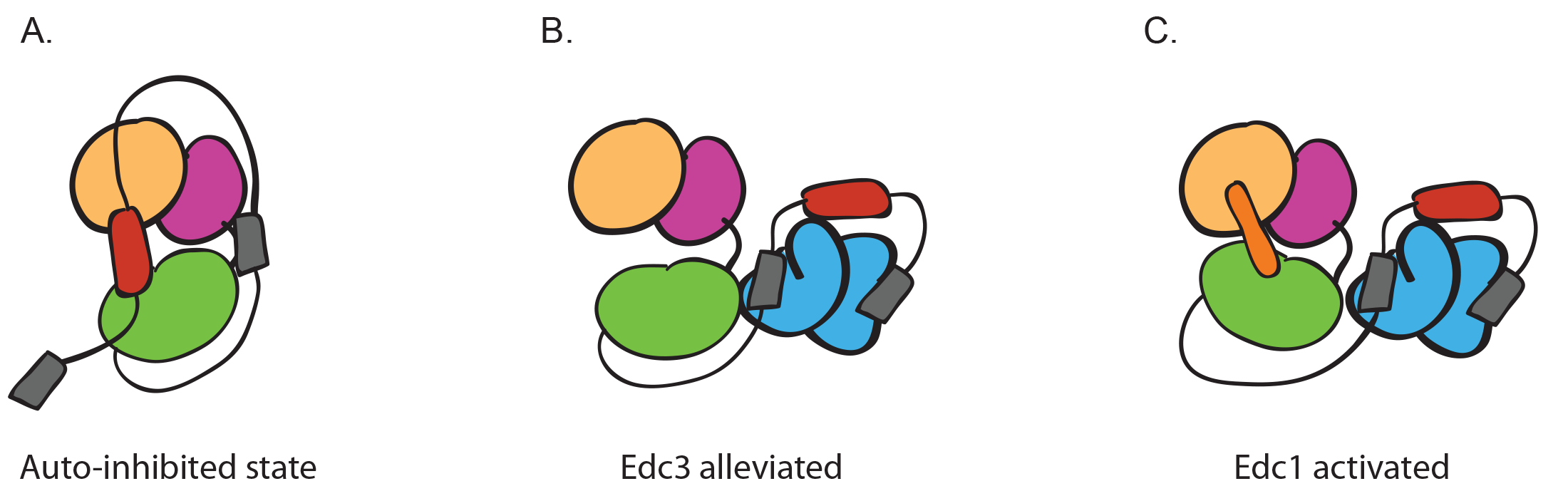
Model for autoinhibition, Edc3 alleviation and Edc3/Edc1 combined activation. (**A**) Autoinhibited conformation of the decapping holoenzyme. Dcp1 is yellow-gold, and the NRD and CD of Dcp2 are magenta and green, respectively. IM1&IM2, shown in red, stabilize this inactive conformation by making contacts with the core domains. The grey boxes are HLMs. (**B**) Edc3, shown in cyan, alleviated inhibition by binding to the HLMs, which disrupts the IM1&IM2 interaction with the Dcp1:Dcp2_core_. (**C**) Representation of the activated Dcp1:Dcp2 structure where Edc1, orange, stabilizes a composite active site formed by the NRD and CD. Edc3 frees up the Dcp1:Edc1 binding site from IM1 & IM2.

In budding yeast, there is clear evidence of autoinhibition, as excision of a short linear motif in the C-terminus bypasses the requirement for Edc3 for degradation of target transcripts (29). Notably, introduction of the equivalent Y220 mutation in budding yeast resulted in a temperature sensitive phenotype, suggesting the cap-occluded conformation may be important for maintaining proper mRNA degradation in vivo (45). The autoinhibitory linear motif identified in budding yeast Dcp2 has similarity to IM1 of fission yeast, whereas IM2 appears to be unique to fission yeast (**Fig S2A,B**). Future studies will be required to identify Edc3 dependent mRNAs in fission yeast and how they are dependent on au toinhibition as described here.

Dcp1:Dcp2_core_ makes fast excursions between the open and cap-occluded states (19, 37), so it is likely the inhibitory motifs in the C-terminal extension stabilize the closed, cap-occluded form by direct interactions with the structured core domains. In support of this hypothesis, the inhibitory motifs found in the C-terminal extension of Dcp2 bind the structured domains of Dcp1:Dcp2_core_ in trans (**Fig S2C** and 37). We cannot exclude the possibility that the inhibitory motifs may stabilize another conformation of the enzyme that is catalytically impaired, such as the open form; or inhibitory motifs could sterically block interactions with substrate. These modes of autoinhibition are reminiscent of the eukaryotic tyrosine kinase superfamily, where motifs flanking the functional domains stabilize an inactive conformation of the enzyme, with structural variation amongst family members (46). Detailed structural studies will be required to define the precise mode of autoinhibition.

How does Edc3 alleviate autoinhibition and allow full activation by Edc1? The Lsm domain of Edc3 is sufficient to alleviate autoinhibition in Dcp1:Dcp2_ext_ and it binds the HLMs found in the Dcp2 C-terminal extension (**Fig 8B**). Prior NMR studies indicate Edc1 and IM1 of the Dcp2 C-terminal extension bind the same surface on Dcp1 (37); moreover, we show here that IM2 binds Dcp1:Dcp2_core_ in trans and that Edc3 can lower the threshold for activation by Edc1. A working model for activation of decapping by Edc3 is that binding to HLMs found in the Dcp2 C-terminal extension is coupled to remodeling of the IM1 and IM2 interaction with the structured, core domains, allowing the enzyme to undergo a conformational change from an inactive to an active, cap-accessible conformation that is stabilized by Edc1 (**Fig 8C**). In this way, the coactivators Edc3 and Edc1 coordinate to destabilize the inactive, autoinhibited form and consolidate the catalytically active conformation, respectively.

Our biochemical and biophysical data are consistent with genetic studies in budding yeast that indicate decapping coactivators work together to promote 5’ mediated decay. First, synthetic growth defects are observed when Edc1 and Edc3 are deleted in yeast strains where Dcp2 is essential (21). Second, deletion of an inhibitory motif in budding yeast Dcp2 bypasses the requirement of Edc3 for decapping on specific RNAs (30). Third, additional proteins such as Scd6 and Pat1 have synthetic genetic interactions with Edc3 and form physical interactions with the HLMs found in the C-terminal extension of Dcp2 (28). Clearly the combinatorial control of Dcp1:Dcp2 activity we observe *in vitro* by Edc1 and Edc3 is well supported by functional studies in cells and the general principles observed here may be applicable to other activators such as Scd6 and Pat1.

In general, any protein that contains an HLM interaction domain such as the LSm domain of Edc3 would promote alleviation of autoinhibition from the Dcp2 C-terminal extension. The fusion of an HLM interaction domain to an RNA binding domain could further increase the activation of the decapping enzyme through increased RNA binding capacity. For example, the C-terminal extension of Dcp2 could provide rationale for Pat1’s strong effect on decapping i*n vivo* (47, 48), since Pat1 can bind HLMs and is linked to the Lsm1-7 complex (49–51) that binds oligoadenylated RNA intermediates during bulk 5’-3’ decay. Further studies will be required to investigate how pathway specific activators such as Pat1 activate decapping through interactions with the C-terminal extension of Dcp2.

We suggest the observations reported in this work on fungal proteins could be conserved in metazoans. The HLM of the fungal Dcp2 has been transferred to a long C-terminal extension of Dcp1 (27). A structure of the Edc3 LSm domain and the HLM in Dcp1 from D. melanogaster is consistent with this interaction being evolutionarily important (20). Transfer of regulatory short linear interaction motifs to different subunits of a conserved complex is a common theme in eukaryotic biology (27, 35). An important area for future research is to determine if the inhibitory motifs of Dcp2 have also been transferred to different subunits of the decapping complex by evolutionary rewiring.

In this work, we have characterized an additional mechanism responsible for regulating mRNA decay by showing that elements in the C-terminus of Dcp2 inhibit decapping at the catalytic step. Addition of Edc3 alleviates autoinhibition to promote catalysis and is required for full stimulation of this extended construct by Edc1. This combinatorial activation suggests decapping by Dcp2 can be controlled across multiple log units of activity to coordinate decay. Such exquisite regulation of decapping reflects the importance of 5’-3’ mRNA decay in maintaining cellular homeostasis.

## ACKNOWLEDGEMENTS

This work was supported by US National Institutes of Health [grant R01GM078360] to J.D.G.; a UCSF Sandler Program for Breakthrough Biomedical Research New Frontier Research award; a National Science Foundation predoctoral fellowship to D.R.P.; and a Genentech Foundation predoctoral fellowship to R.W.T.

## AUTHOR CONTRIBUTIONS

D.R.P. and R.W.T designed and implemented the experiments. T.S.D. implemented experiments. D.R.P., R.W.T. and J.D.G. wrote the paper. J.D.G. supervised the research.

